# Diverse Defence Systems and Prophages in Human-Associated *Bifidobacterium* Species Reveal “Arms Race” Dynamics

**DOI:** 10.1101/2025.01.20.633837

**Authors:** James A. D. Docherty, Ryan Cook, Raymond Kiu, Xena Dyball, Teagan Brown, Magdalena Kujawska, Rhianna lily-Smith, Sarah Phillips, Rachel Watt, Andrea Telatin, Sumeet Tiwari, Lindsay J. Hall, Evelien M. Adriaenssens

**Author notes:** Corresponding author, Quadram Institute Bioscience, Norwich Research Park, Norwich, UK.

## Abstract

Bacteria of the genus *Bifidobacterium* are pivotal for human health, especially in early life, where they dominate the gut microbiome in healthy infants. Bacteriophages, viruses of bacteria, are drivers of gut bacterial composition in the human gut and could affect bifidobacterial abundance. Here, we use a bioinformatics approach to explore the direct interactions occurring between human-associated *Bifidobacterium* spp. and prophages, as evidenced by their genomes. A total of 1,086 bifidobacterial genomes were analysed in this study, revealing complex systems to prevent viral invasion. Despite their characteristically small genomes, *Bifidobacterium* strains harboured more than double the number of defence systems as most bacteria. In total, 34 defence system types and 56 subtypes were detected, including several different CRISPR-Cas systems with spacers that targeted almost three-quarters of bifidobacteria-derived prophages. We identified at least one prophage which met our stringent quality control measures in ∼63% of strains, with phages exhibiting high genomic diversity and evidence of historical recombination. Additionally, prophages were found to encode various anti-defence systems, such as anti-CRISPR genes and restriction modification resistance mechanisms. In summary, our investigation reveals “arms race” dynamics drive genomic diversity in both bifidobacteria and their phages.

**Importance:** Members of the *Bifidobacterium* genus are widely acknowledged as being highly important for human health, particularly in infants. To date, there have been a limited number of large-scale studies that have investigated the presence of prophages and anti-viral defence systems of *Bifidobacterium* strains from multiple human-associated species. Here, we have uncovered a complex set of anti-phage strategies encoded by *Bifidobacterium* strains. In addition, we have also identified a highly diverse phage mobilome present within the genomes of bifidobacteria across the genus, which also encode several unique systems for overcoming bacterial defences. Elucidating these co-evolutionary dynamics between phages and bifidobacteria may provide valuable insight into developing high-throughput methods for identifying next-generation probiotic or live biotherapeutic candidates, which may then be applied for the prevention and treatment of various diseases in humans.

## Introduction

Members of the *Bifidobacterium* genus are key members of the human gut microbiome, where they are one of the earliest and most important colonisers of the human gut (1). In healthy breast-fed infants, bifidobacteria are highly abundant, and can comprise ≥70% of all microbial cells present in the gastrointestinal tract (2, 3, 4). Reduction in their abundance at this critical stage is correlated with numerous disorders, including allergies (5), autoimmune diseases (6) and necrotising enterocolitis (7, 8). Despite the levels of *Bifidobacterium* species being lower in adults, with a relative abundance of 2-14% (9), their presence is still central to host health, as reductions are associated with a range of diseases, including inflammatory bowel disease (10, 11) and colorectal cancer (12, 13).

While reduced levels of bifidobacteria and gut microbiota perturbations are known to be associated with a variety of factors, such as birth mode (14), age (9, 15, 16), diet (17, 18) and antibiotic use (19), a potentially overlooked modulator of the gut ecosystem are bacteriophages. While phages are present in the human gastrointestinal tract at lower proportions relative to bacteria (∼1:1) (20, 21, 22) compared to alternative niches, such as aquatic environments and soil (∼10:1) (23, 24, 25, 26, 27, 28), they are thought to drive shifts in bacterial composition through various ecological models, including arms race dynamics (29, 30). This concept implies that phage infection exerts a directional selection pressure on the host, which favours the development of resistant populations. In turn, this drives mutations in the phage to overcome these defences and restore infectivity, prompting an ongoing cycle of resistance and counter-resistance (30, 31).

Reports have suggested that the induction of prophages in bifidobacteria may reduce their abundance in the gut microbiome of infants (32). This observation may be attributed to a process known as “community shuffling”, which has the potential to cause disturbances by decreasing the abundance of beneficial bacteria, thereby allowing reoccupation of the niche by microbiota members with pathogenic potential and pathogens (33, 34, 35). To combat the persistent threat from phages, bacteria have evolved a diverse and effective defensive arsenal targeting their viral invaders, which traditionally, were primarily thought to be comprised of Restriction Modification (RM), Abortive Infection (Abi), and more recently, CRISPR-Cas systems (36, 37, 38). However, a plethora of bacterial “immune systems” have newly been uncovered, with more than 200 anti-viral defence systems described to date (36, 39, 40, 41). These discoveries have led to the development of new bioinformatics tools for rapid and accurate prediction of these defence systems in bacterial genomes (40, 42), which has opened new avenues for the exploration of the intricate relationships between bacteria and phages. It has also recently been confirmed that defence systems display synergism to enhance protection from phages (43).

Although there has been extensive *Bifidobacterium* research, the functional annotation of their genomes remains rather cryptic. Their highly specialised genomes are characteristically small, generally ranging from just 1.6-3.2Mbp (44), compared to the “typical” ∼5Mbp for most bacteria (45), such as *Escherichia coli* (46). Despite ongoing efforts to decipher functional annotations, many of their gene functions remain unknown (47, 48). In addition, most existing literature has focussed on investigating genetic differences each within a single different *Bifidobacterium* species (49, 50, 51, 52, 53), with relatively few reports detailing inter-species differences (54, 55). However, patterns have emerged from these individual studies, such as a high occurrence of CRISPR-Cas systems and prophages in several *Bifidobacterium* species, including important probiotic candidates such as *B. pseudocatenulatum* (49, 56) and *B. longum* (57). The antiviral arsenal of the Actinomycetota (formally Actinobacteria) phyla has also recently been considered, which included 120 *Bifidobacterium* genomes from the RefSeq database (58). Using a bioinformatics-based approach, we aimed to investigate the extent to which human-associated bifidobacteria and phages contribute to each other’s genomic diversity through antagonistic interactions. Here, we propose that dynamic phage-host interactions are driving bifidobacteria to continuously adapt their defence systems to fend off viral invaders, while phages are evolving counterstrategies to circumvent these defences.

## Methods

### Ethics statement

The Pregnancy and EARly Life (PEARL) study (59) was approved by the Human Research Governance Committee at QIB and the London-Dulwich Research Ethics Committee (reference 18/LO/1703). Written ethical approval was received from the Human Research Authority (IRAS project ID number 241880). Faecal samples from healthy British born infants (full-term, breast-fed) were collected using methods compliant with protocols established by the National Research Ethics Service-approved UEA/QIB Biorepository (License no: 11208) and the Quadram Institute Bioscience Ethics Committee (60).

### Isolation of *Bifidobacterium* strains from PEARL samples and British infants

Maternal stool samples (PEARL study) (59), which were previously collected and frozen at set time points, were thawed and mixed with phosphate buffered saline (PBS) (100mg of sample with 1ml of PBS). The resulting homogenised faecal slurry was then serially diluted to 10^3^, with 200μl of dilutions 10^2^ and 10^3^ then being plated onto selective Brain Heart Infusion (BHI) agar plates containing 1ml of cysteine and mupirocin per 1L of media. Following incubation at 37°C in an anaerobic chamber for a maximum of 48 hours, five colonies that resembled the morphology of *Bifidobacterium* strains were selected for quadrant streaking on selective agar for a further 48 hours. Isolates were re-streaked three times to purity. Frozen stocks of pure culture were prepared and stored in cryovials at –80°C (30% glycerol in BHI media).

For *Bifidobacterium* strains isolated from healthy British born infants (60), faecal samples were initially streaked onto Reinforced Clostridial Medium (RCM) agar supplemented with cysteine and mupirocin (0.05mg/ml). Colonies were then randomly selected and re-streaked onto de Man Rogosa and Sharpe agar plates supplemented with 0.05mg/ml mupirocin and 500mg/ml cysteine to purity. Frozen stocks of pure culture were prepared and stored in cryovials at –80°C (30% glycerol in RCM media).

### Genomic DNA extraction and whole genome sequencing of bacterial isolates

Genomic DNA of bacterial isolates from the PEARL study were isolated in accordance with the protocol provided by the Maxwell® RSC PureFood GMO and Authentication Kit. The Maxwell® RSC was prepared and run in compliance with the PureFood Protocol. Whole genome sequencing was performed using the Illumina NextSeq500 platform with read lengths of 125 bp (paired end reads) by Quadram Institute Bioscience (QIB) Sequencing Core Services. For information regarding the DNA extraction and sequencing of genomes from British infants, please see the ‘Methods’ section from Lawson *et al*. (60).

### Datasets used for downstream analysis

All publicly available *Bifidobacterium* genomes were downloaded from the National Library of Medicine (National Center for Biotechnology Information) (https://www.ncbi.nlm.nih.gov/datasets/genome/?taxon=1678) in July 2023 from the GenBank database (61). The search option ‘Exclude atypical genomes’ was used prior to downloading assemblies (https://www.ncbi.nlm.nih.gov/assembly/help/anomnotrefseq/).

FastQ files for 48 *B. pseudocatenulatum* strains from project PRJNA720750 (study title: ‘Exploring the Genomic Diversity and Antimicrobial Susceptibility of *Bifidobacterium pseudocatenulatum* in a Vietnamese Population’) (62) were downloaded from the European Nucleotide Archive (https://www.ebi.ac.uk/ena/browser/home). The quality control of raw-reads and followed by assembly was performed using the Galaxy platform, hosted by QIB. (https://f1000research.com/slides/8-1663 and https://doi.org/10.1093/nar/gkae410).

FastQC (v0.72) was used under default parameters to determine the read quality of raw sequence data (63). Pre-processing of the raw sequence data, including global and adaptor trimming, was performed using fastp (v0.23.2) (64). Trimmed reads were then assembled using Shovill with default settings (v1.1.0) (65, 66).

Prior to quality checks (QC) and dereplication of bacterial genomes at strain level (see below), there were 28 genomes for strains isolated from healthy British born infants were included in this study (BioProject accession no. PRJEB28188) (60), as well as 22 strains from Cambodian infants (BioProject accession no. PRJNA958097) (67). There were also 133 assemblies for strains isolated from participants of the PEARL study included in this investigation (59).

A full list of *Bifidobacterium* genomes used in the final database of this study, including accompanying metadata, is available in Table S1 in supplemental material. Metadata was visualised using the Microreact web application (https://microreact.org/) (68).

### Quality checks and dereplication of human-associated *Bifidobacterium* genomes

To select for genomes exclusively isolated from humans (NCBI dataset), a Python script was used that would accept and search for a query string in individual lines (“*Homo sapiens*”) from the assembly data report (see supporting code). All genome assemblies were quality-checked using CheckM (v1.2.0) (69). The typical workflow option ‘*lineage_wf*’ was used to determine genome quality and contamination. Only genomes of high quality (≥90% completeness, ≤5% contamination) were retained. Next, dRep (v3.4.3) was used to dereplicate genomes at strain level (average nucleotide identity (ANI) cut-off of ≥99.9%) (70). Following dereplication, there were a total of 1,244 genomes. As a final layer of QC, the SeqFu (1.20.3) (71) subprogram ‘*seqfu stats*’ was used to obtain assembly statistics for each genome in the dereplicated dataset. All genomes with an N50 below 50,000 or comprised of more than 100 contigs were removed from the final database.

A total 1,086 genomes from 15 *Bifidobacterium* species, as well as a single *Gardnerella vaginalis* strain (previously *Bifidobacterium piotii*) included as an outgroup, formed the bacterial database for downstream analysis.

### In silico analysis of Bifidobacterium genomes

All genomes in the filtered database underwent genome annotation using Prokka (v1.14.6) (default: ‘--centre, –-compliant’) (72) to generate a protein sequence file (.faa) for each genome. PhyloPhlAn (v3.0.67) (parameters: ‘--fast, –-verbose, –-supertree_aa.cfg’) (73, 74) was used to construct a phylogenetic tree for all strains. The tree visualisation was performed using iTol (v5) (75). Further taxonomic assignment of *Bifidobacterium* genomes was performed using GTDB-Tk (v2.1.0) (76, 77), workflow option ‘*classify_wf*’. Strain classifications were adjusted accordingly based on discrepancies identified in their phylogenetic classification. See Table S2 in supplemental material for any changes made to species names.

Using the amino acid sequence and general feature format files generated by Prokka as inputs, the presence of anti-viral defence systems in bacterial genomes was predicted using DefenseFinder (v1.3.0) (40) and PADLOC (Prokaryotic Antiviral Defence LOCator) (v2.0.0) (78). Defence systems categorised as ‘*adaptation*’ and ‘*other*’ by PADLOC were discarded from the analysis to ensure that only well characterised systems were included in the final counts. Additionally, all defence systems labelled as a ‘phage defence candidate’ (PDC) or ‘Hma-embedded candidate’ (HEC) were excluded, so that only validated systems were accepted. As part of the final concatenation and deduplication of results from each tool, the output from PADLOC was first processed to match/align with the structure of the DefenseFinder output files (see supporting code). Due to differences in naming conventions between DefenseFinder and PADLOC, as well as variation in the calling of defence system types, overlaps were manually curated and amended (see Table S3).

Coinfinder (v1.2.1) (79) was employed to detect positively co-occurring and dissociating pairs of defense systems within bifidobacterial genomes. Additionally, the Newick tree file generated by PhyloPhlAn (73) was also used as input. A network of associating and dissociating genes was visualised using Cytoscape (v3.10.1) (80) and annotated using Adobe Photoshop (v25.0) (81). The localisation of co-occurring defence systems was further investigated, whereby defence system pairs were considered localised when ≥50% of occurrences were within 20 genes.

CRISPRCasTyper (v1.8.0) (82) was used to predict CRISPR arrays in bacterial genomes. Unique spacers were identified following dereplication by CD-HIT (v4.8.1) (83) with a sequence identity threshold of 100%. Sequence matches between spacers and prophages were predicted using the ‘*predictmatch*’ utility in SpacePHARER (v5.c2e680a) (84).

### Identification of *Bifidobacterium* prophages

Prophage regions were identified within bifidobacterial genomes *in silico* using the computational tools, VIBRANT (v1.2.1) (85), geNomad (v1.6.1) (86) and PhageBoost (0.1.7) (87). Each tool was run under default parameters. Overlapping prophage coordinates from each tool were merged to include the earliest start and the latest end coordinate within the host bacterial scaffold. Prophage regions were then located and excised from their respective cognate host genome (see supporting code).

The quality and completeness of all initial predictions were assessed using CheckV (v1.0.1) (88), and trimming of bacterial contamination on the prophage ends was performed where indicated by the program output. All contigs assigned as “Low-Quality” and fragments under 5kb were immediately discarded. Manual curation of the ten largest prophages categorised as “Not-Determined”, which all had no viral genes detected using CheckV, was conducted.

Putative prophages from this category were then excluded from the database. Only prophages categorised as “Medium-Quality” or higher by CheckV, meaning a predicted genome completeness over 50%, were then further scrutinised prior to finalising the prophage database.

Using Pharokka (v1.7.1) (89), putative prophage sequences were annotated under default parameters. This tool utilises PHANOTATE to predict coding sequences (CDS) (90) and tRNAscan-SE 2.0 (91), Aragorn (92) and CRT (93) for the prediction of tRNAs, tmRNAs and CRISPRs, respectively. Functional annotation was achieved by matching CDS to the PHROGs (94), VFDB (95) and CARD (96) databases using MMseqs2 (97) and PyHMMER (98). The INPHARED database (version as of April, 2024) (99) was also utilised to match contigs to their closest hit using Mash (100). Plots were generated using the pyCirclize tool (101). Only phages containing at least four structural proteins (categorised as ‘Head and Packaging’, ‘Tail’ and ‘Connector’) and comprising ≥10% of CDS (based on nine functional categories, as assigned by PHROGS (94)) were accepted for the final prophage database.

### Preliminary characterisation of bifidobacteria-derived prophages

Prophage genomes were then dereplicated at species level via genome clustering based on pairwise ANI (≥95% ANI cutoff & ≥85% target coverage; MIUViG recommended-parameters) using BLAST (v2.13.0) alongside supporting code from CheckV (https://bitbucket.org/berkeleylab/checkv/src/master/) (88, 102, 103). Prior to the completion of the dereplicated prophage database, intergenomic similarity of all prophages, based on pairwise ANI (from the CheckV supporting code), was calculated as a percentage and converted into a matrix. The sum of the query and target sequence values were halved to establish reciprocal intergenomic similarity. A two-way hierarchical clustering heatmap was produced using Heatmap3 (v1.1.9) (104) in R (v4.3.1).

To construct a phylogenetic tree for recovered prophages, all sequences were processed with ViPTreeGen (v1.1.2) to generate a phage proteomic tree (105). Then, for tentative taxonomic classification of prophages into prospective Order ranks, the most up-to-date bacteriophage isolate complete genome database (RefSeq, as of August 2, 2024) was downloaded from INPHARED (https://github.com/RyanCook94/inphared) (99). Then, all phage genomes, including dereplicated *Bifidobacterium* prophages identified in this study, were once again processed with ViPTree. The phylogenetic trees were visualised and edited using iTol (v5) (75). TaxmyPHAGE (v0.2.9) was used in attempts to assign genus level taxonomy (https://github.com/amillard/tax_myPHAGE). The ‘*annotate*’ module from geNomad (86) was used to determine if each of bifidobacteria-derived prophages belonged to the *Caudoviricetes* class.

### Identification of “anti-defence” genes in prophages

Anti-CRISPR (*acr*) and anti-CRISPR-associated (*aca*) genes in prophages were identified using Acafinder (version as of November 1, 2023; ‘-w/--Virus optionߣ, guilt-by-association approach) (106). Supporting code from dbAPIS (a database of anti-prokaryotic immune system genes) (https://github.com/azureycy/dbAPIS) (107) was used to identify alternative anti-prokaryotic immune system proteins via an HMMscan of whole-genome protein sequences (.faa files).

### Data Analysis and Visualisation

All statistical analyses were performed using base R statistical package (v4.3.1) and graphs were visualised using R package *ggplot2* (v3.5.1) (108) in RStudio (v2023.6.2.561) (109), unless stated otherwise.

### Supporting code

Supporting code is available at https://sweet-diadem-981.notion.site/Diverse-Defence-Systems-and-Prophages-in-Human-Associated-Bifidobacterium-Reveal-Arms-Race-Dynamic-14bd61222c0380538849facd06996ed9.

## Results

### *Bifidobacterium* spp. encode a diverse repertoire of antiviral defence systems

After quality control and strain level dereplication, a total of 1,087 complete and draft genomes from at least 29 countries were used to investigate the antiviral defence systems and prophage repertoire of human-associated bifidobacteria (Figure S1). Taxonomic classification of strains revealed that there were 15 individual *Bifidobacterium* species and a single *Gardnerella* strain which we retained as an outgroup (Figure 1). The final bifidobacterial genome database was comprised of *Bifidobacterium adolescentis* (n = 171), *Bifidobacterium angulatum* (n = 2), *Bifidobacterium animalis* (n = 2) (subsp. *animalis*, n = 1; subsp. *lactis*, n = 1), *Bifidobacterium bifidum* (n = 82), *Bifidobacterium breve* (n = 91), *Bifidobacterium catenulatum* (n = 38) (subsp. *catenulatum*, n = 1; subsp. *kashiwanohense*, n = 21), *Bifidobacterium dentium* (n = 19), *Bifidobacterium gallicum* (n = 1), *Bifidobacterium longum* (n = 473) (subsp. *longum*, n = 160; subsp. *infantis*, n = 20; subsp. *suis*, n = 2), *Bifidobacterium pseudocatenulatum* (n = 196), *Bifidobacterium pseudolongum* (n = 7) (subsp. *globosum*, n = 1), *Bifidobacterium pullorum* (n = 1) *Bifidobacterium ruminantium* (n = 1), *Bifidobacterium scardovii* (n = 1), *Bifidobacterium thermophilum* strain (n = 1) and one *Gardnerella vaginalis* strain. After profiling, there were several discrepancies identified between the labelling of species names and their phylogenetic classification, which were corrected based on the phylogenomic profile and GTDB taxonomy assignment (Table S2).

**Figure 1.**
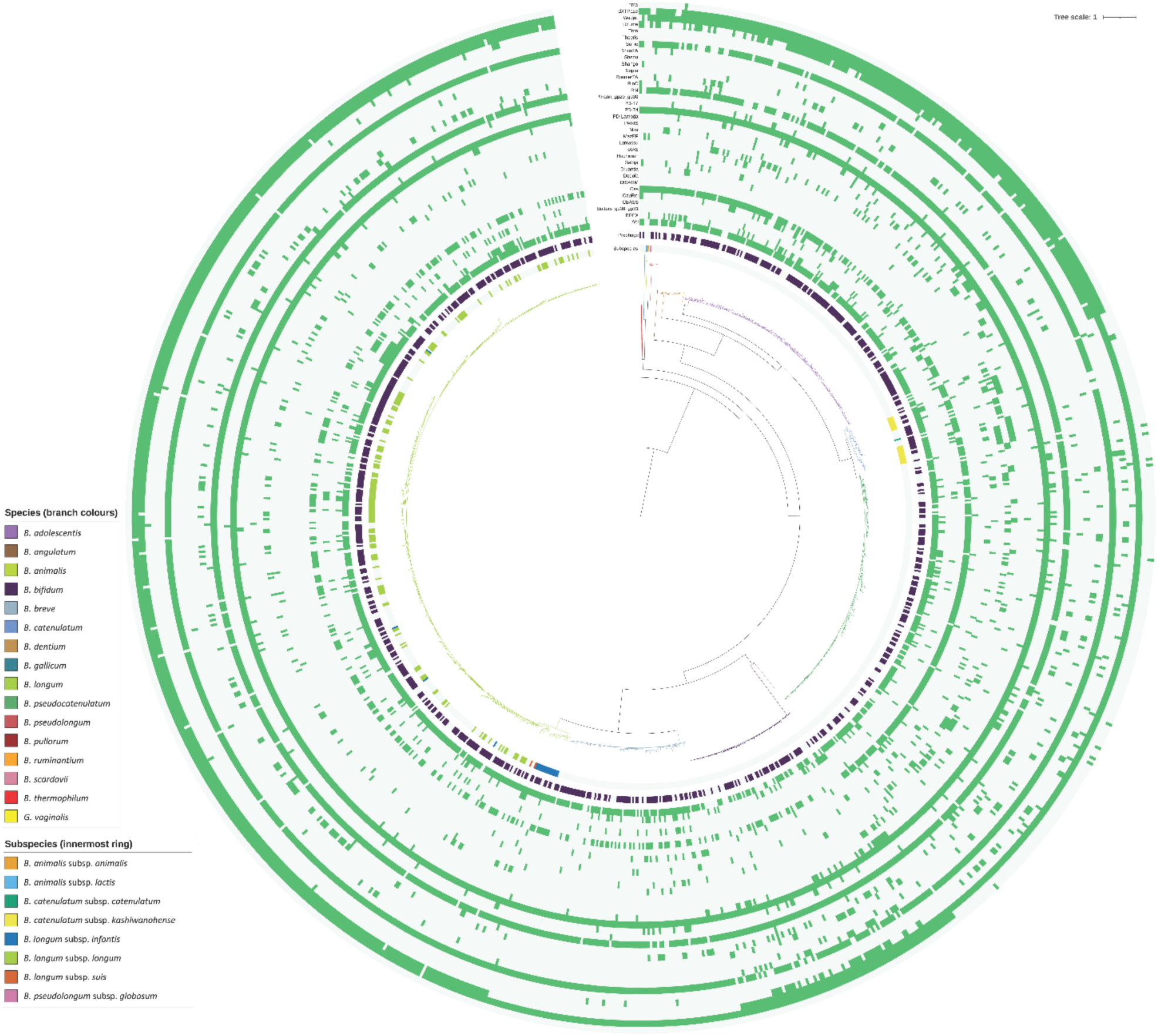
Phylogenetic tree of Bifidobacterium genomes, based on 400 universal bacterial ‘marker’ genes. Branch colours represent individual species, and the innermost ring is coloured in accordance with the subspecies designation where available. The second ring corresponds to the presence (purple) or absence (light grey) of predicted prophage sequences that are more than 50% complete in bifidobacterial genomes. The outer heatmap represents the presence (green) or absence (light grey) of individual defence systems, grouped per defence system type. The tree scale and branch lengths correspond to the number of substitutions per site. From the innermost ring of the outer heatmap, the defence system types identified in at least one genome were: Abi, BREX, Butters_gp30_gp31, CBASS, CapRel, Cas, DISARM, Dodola, Druantia, Gabija, Hachiman, IetAS, Lamassu, MazEF, Mza, PARIS, PD-Lambda, PD-T4, PD-T7, Phrann_gp29_gp30, RM, RloC, RosmerTA, Septu, Shango, ShosTA, SoFic, Thoeris, Tmn, Uzume, Wadjet, dXTPase and hma.

Deduplication of defence systems revealed that there was a total of 13,185 anti-viral mechanisms from 1,086 *Bifidobacterium* genomes, as well as a single *G. vaginalis* strain, showing a mean of 12.13 and a median of 12 defence systems per genome. The number of defence systems for each genome was found to demonstrate considerable variation across individual strains (±SD = 2.67), ranging from as few as 5 systems to as many as 25. Further inspection of systems revealed that there were at least 34 defence system types and 56 subtypes that had a varied distribution across the dataset (Figure 1), each with varying percentage occurrences (Figure S2; Table 1). At least one PD-T4 defence system was identified in every human-associated *Bifidobacterium* genome, with a further three defence system types occurring in ≥90% of strains, namely: RM (94.57%), SoFic (94.39%) and Wadjet (90.24%). The next most prevalent systems were Abi (83.26%), dXTPases (78.01%) and CRISPR-Cas (58.51%). There were also 16 systems that occurred in fewer than 5% of strains, such as Gabija and RloC, which occurred in 4.88% and 3.40% of strains, respectively.

**Table 1.** Percentage occurrence of defence system types in human-associated *Bifidobacterium* genomes. List of defence systems and their percentage occurrence in bifidobacteria in descending order.

| Defence System | Abbreviation | Occurrence (%) |
| --- | --- | --- |
| Phage Defence Type 4 | PD-T4 | 100 |
| Standalone protein with Fic domain | SoFic | 94.39 |
| Restriction Modification | RM | 94.11 |
| Wadjet | Wadjet | 90.25 |
| Abortive Infection | Abi | 83.26 |
| dXTPase | dXTPase | 78.01 |
| CRISPR-Cas | Cas | 58.51 |
| Bacteriophage Exclusion | BREX | 19.87 |
| Uzume | Uzume | 19.60 |
| Lamassu | Lamassu | 16.10 |
| RosmerTA<br>(toxin-antitoxin) | RosmerTA | 15.82 |
| Phage Defence Lambda | PD-Lambda | 9.75 |
| IetAS | IetAS | 9.20 |
| Cyclic oligonucleotide-based<br>antiphage signaling system | CBASS | 9.20 |
| Phage Defence Type 7 | PD-T7 | 6.72 |
| Shedu | Shedu | 6.16 |
| Septu | Septu | 5.61 |
| CapRel | CapRel | 4.97 |
| Gabija | Gabija | 4.88 |
| Mza | Mza | 4.32 |
| Tmn | Tmn | 4.23 |
| RloC | RloC | 3.40 |
| MazEF | MazEF | 2.67 |
| Druantia | Druantia | 2.39 |
| DISARM | DISARM | 2.30 |
| ShosTA<br>(toxin-antitoxin) | ShosTA | 2.21 |
| Dodola | Dodola | 1.56 |
| PARIS | PARIS | 1.20 |
| hma | hma | 1.01 |
| Butters_gp30_gp31 | gp30_gp31 | 0.55 |
| Phrann_gp29_gp30 | gp29_gp30 | 0.55 |
| Thoeris | Thoeris | 0.46 |
| Hachiman | Hachiman | 0.28 |

The prevalence of defence system types across individual species was inspected to identify inter-specific differences in their distribution (Figure S3). Notable trends were immediately apparent, Wadjet was highly prevalent (∼98.5%) in every *Bifidobacterium* species (>5 strain representative genomes), except *B. breve*, where the defence system was completely absent. Additionally, there were also substantial differences in the presence of CRISPR-Cas systems between individual species. For example, there were Cas systems present in just 34.15% of *B. bifidum* strains and 94.74% of *B. dentium* strains. Furthermore, the defence system Uzume was detected in ∼80% of *B. adolesecentis* genomes. However, it was completely absent in several other bifidobacterial species, including *B. breve* and *B. catenulatum*, and present in less than 10% of *B. longum* and *B. pseudocatenulatum* genomes. *B. pseudocatenulatum* strains didn’t encode a single dXTPase system, while it was found in over 78% of all strains, including all *B. bifidum*, *B. breve*, *B. dentium* genomes, as well as in more than 99% of *B. adolescentis* and *B. longum* genomes.

To further explore the distribution of defence systems across the genus, the average number of defence systems within each species was also examined. It was discovered that there were significant differences observed between the number of defence systems (p ≤ 0.05, Kruskal-Wallis’ test), in addition to significant differences between several independent species pairs (p ≤ 0.05, Dunn’s test) (Figure 2). The median number of defence systems ranged from 10.2 in *B. dentium* to 12.8 in *B. catenulatum* (>5 strains analysed). Assessment of the distribution of defence system types across individual *Bifidobacterium* genomes indicated that there was no significant association between single strains (Chi-square statistic with Monte Carlo simulation of 31,674.68 and a simulated p-value of 1; 10,000 simulations). Though, there was a significant difference in the presence of defence system types between different *Bifidobacterium* species (Chi-square statistic with Monte Carlo simulation of 4,133.372 and a simulated p-value of <0.05; 10,000 simulations).

**Figure 2.**
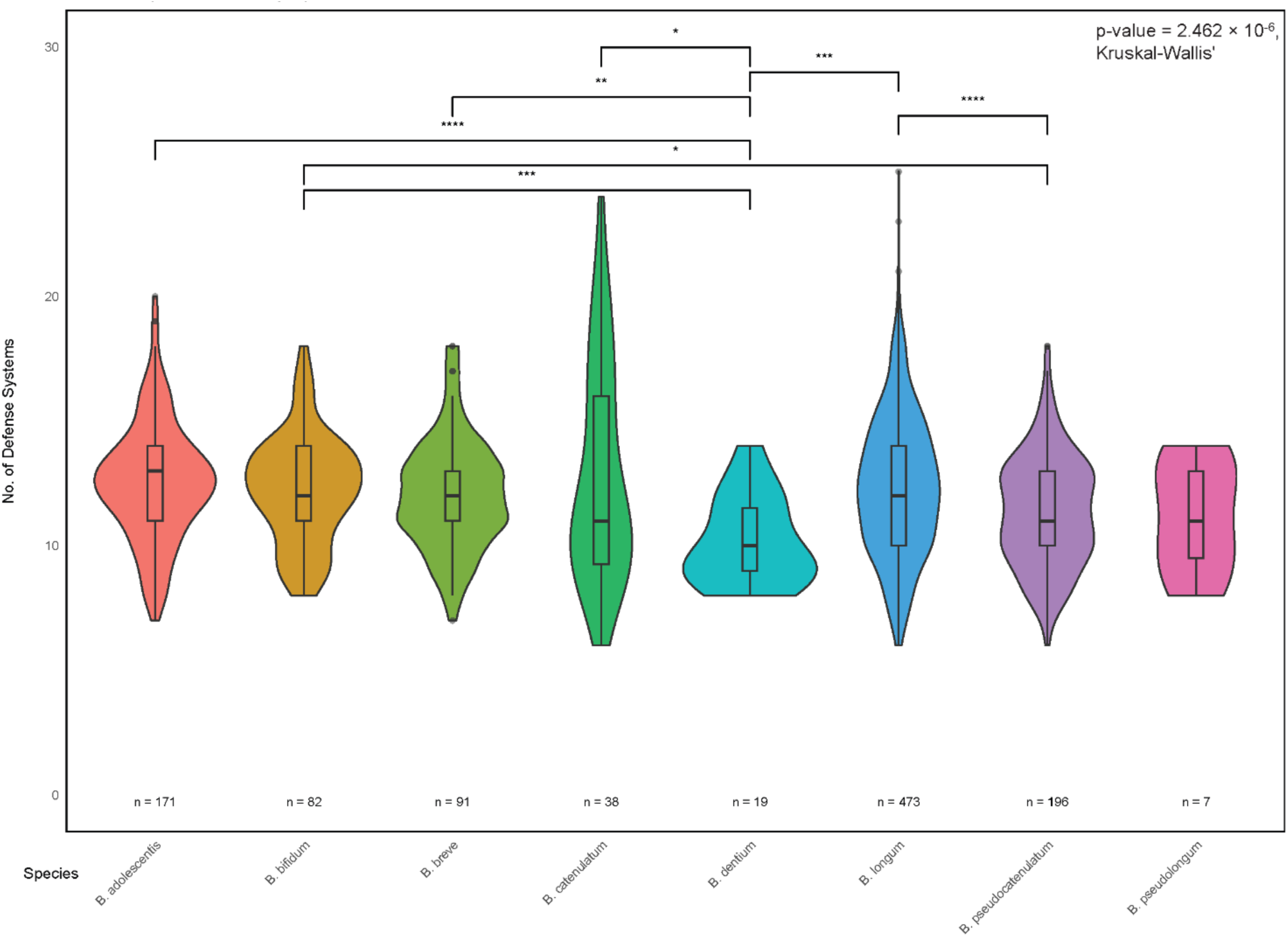
Violin plots depicting the median of defence systems in individual species (>5 representative strains). Kruskal-Wallis chi-squared = p-value < 0.05. Colours represent individual species and the number of strains corresponds to ‘n = X’. Dunn’s test significance is indicated where *: p-value < 0.05, **: p-value < 0.01, ***: p-value < 0.001, ****: p-value < 0.0001.

In summary, our results revealed that bifidobacteria encode a broad range of different defence systems, with certain mechanisms occurring frequently among several species across the genus and others present in just a few strains, such as Gabija and RloC.

### Several pairs of defence systems positively co-occur in bifidobacteria

Here, we aimed to investigate if pairs of defence systems occurred together more or less frequently than would be expected by chance, i.e. whether they associate or dissociate. Given the close phylogenetic ties within bifidobacteria, the co-occurrence analysis was performed with consideration of evolutionary distances from the phylogenetic analysis. In total, there were 21 pairs of positively associating defence systems and 11 pairs of dissociating systems out of 428 pairs tested (Bonferroni significance correction = p-value < 0.01) (Figure 3). There were six instances where CRISPR-Cas systems were associated with different RM and Abi systems. CRISPR-Cas Type I-C was found to associate with two Abi systems, namely Abi2 and AbiD. CRISPR-Cas Type II-C and I-G were also found to positively co-occur with AbiE and AbiL, respectively. Type II-C and I-E Cas systems were also found to co-occur with RM Type II-G and Type-IV. There were multiple cases where several Abi systems separately co-occurred with other abortive infection/growth inhibitors, such as CBASS_I, RosmerTA and ietAS. CBASS_I was also discovered to associate with BREX_I, with BREX_I also co-occurring with Shedu. Of the 11 pairs of dissociating pairs of defence systems, more than half of these pairs included dXTPase, which was found to predominantly negatively associate with abortive infection-type systems, including Abi2, AbiD, ietAS and mechanisms belonging to the Lamassu-Family. The dXTPase was also found to dissociate with RM Type-IIG and CRISPR-Cas Type I-C. Cas Type I-C was also found to dissociate with AbiE and RM Type-IV.

**Figure 3.**
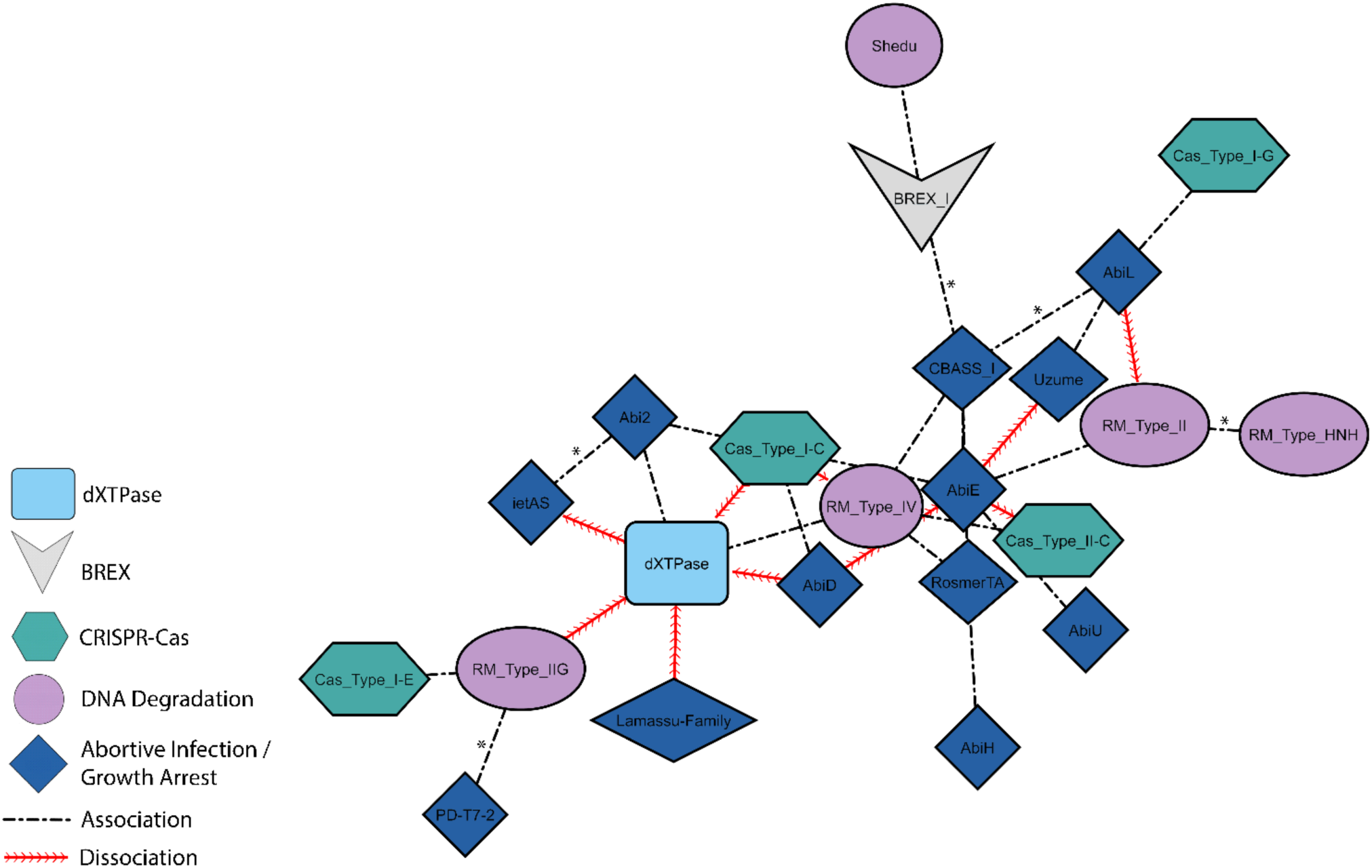
Network of co-occurring and dissociating pairs of genes in bifidobacteria. Network visualisation of statistically significant (Bonferroni significance correction = p-value < 0.01) positively co-occurring (straight lines) and dissociating pairs (sinewave) of defence system subtypes in bifidobacteria. The length of connecting lines is proportional to the strength of the relationship (significance score). An asterisk (*) denotes a pair of systems that are co-localised.

Of the 21 pairs of positively co-occurring defence systems found in human-associated bifidobacteria, just six were found to co-localise. Namely, the localised pairs of defence systems were: BREX I with CBASS I and Shedu, AbiL with CBASS I, Abi2 with IetAS, PD-T7-2 with RM Type II-G and RM Type HNH with RM Type II.

### Adaptive immunity in human-associated bifidobacteria

Exploration into the adaptive immunity repertoire of human-associated bifidobacteria revealed that 58.51% of strains encoded CRISPR-Cas systems, with at least six unique systems having been identified. The CRISPR-Cas subtypes were inclusive of three Type-I systems (I-C, I-E and I-G), two Type-II systems (II-A and II-C) and one Type-V system, as well as undetermined Cas clusters. The most abundant was CRISPR-Cas I-C, with 232 individual systems identified. There were also three systems where the counts were more than 100, including Type II-C (125), I-E (119) and I-G (109). In addition, there were 71 Type II-A systems and just two Type-V systems, as well as 12 unidentified Cas clusters. With regards to Type II-C systems, in several instances, the *cas2* gene appeared to be degraded/absent from the system, though there were RecD enzymes present, as well as a DEDDh exonuclease. In various bacterial species, DEDDh family exonucleases have been found to be fused with degraded *cas*2 genes (110, 111), suggesting that the lost nuclease activity of the Cas protein can be restored through an unrelated enzyme (112). The presence of the RecD enzyme could facilitate the acquisition of novel spacers, aiding adaptation, which has been observed in *E. coli* (113). CRISPR-Cas systems were predominantly found to occur on their own in *Bifidobacterium* genomes, though two or more systems did occur together multiple times (Figure S4). The I-E and II-C co-occurred in 18 genomes, as well as I-G and II-C in six cases. I-C and II-C occurred together three times, with I-C and I-E occurring together just twice.

To investigate the specific targets of the CRISPR-Cas systems, CRISPR spacers were tested for their sequence identity with the bifidobacteria-derived prophages. Initially, 30,489 spacers were identified in ∼62% (671/1,087) *Bifidobacterium* genomes with a median of 41 spacers per genome. Following dereplication of spacer sequences at 100% identity, there were 15,809 non-redundant sequences. These dereplicated spacer sequences were found to target ∼74% (798/1,076) of all prophages identified in host strains (Table S4). Further inspection of the interactions of dereplicated spacers with prophages revealed that there were at least 434 host genomes that contained CRISPR sequences that matched proviruses. Overall, there were 7,877 independent spacer-phage interactions, encompassing 4,317 unique spacers, with single spacers found to target multiple phages, as well as single phages being targeted by multiple spacers. Of the 798 phages targeted by the CRISPR spacers, just 151 prophages were targeted by a single spacer. There were 26 phages that shared sequence identity with at least 50 spacers, and three phages with more than 100 hits.

Spacer sequences belonging to the CRISPR-Cas subtype I-C dominated the dataset, with almost 5,000 spacer hits to phages coming from this system. There were more than 1,500 spacer matches from subtype I-E and over 1,200 from subtype I-G. Type II systems comprised less than 300 of total matches with prophages. Investigations into self-targeting spacers demonstrated that just 16 spacers targeted a prophage within the same strain as the host. There was also strong evidence to suggest that there was phage infection occurring across species boundaries, as almost half of all the spacers targeted prophages that were detected in different *Bifidobacterium* species. Additionally, of the 434 *Bifidobacterium* genomes harbouring spacers, 176 of these strains (∼41%) did not contain a prophage (as determined by the prophage detection pipeline), accounting for 3,726 spacers.

### Prophages are genomically diverse and abundant in bifidobacteria

A consolidated prophage recovery workflow was employed to computationally predict proviral regions within the bifidobacterial genomes. All prophages that were labelled as “Medium-quality” or higher (≥50% completeness prediction) by CheckV (88) and with a fragment size over 5kb were further scrutinised for the presence of ≥4 structural proteins comprising at least 10% of CDS. After filtering, there were 1,075 prophages from 732/1,086 (∼67%) *Bifidobacterium* genomes, revealing a mean of 0.99 phages present per host genome, which were included in the final prophage database. These prophage genomes and partial genomes demonstrated a broad size range, varying from 8,930bp to 65,279bp, with the median genome length of 34,740bp. Direct comparison of the GC content of phages and their hosts showed that there was a significant difference between the two values (p < 0.05, Mann-Whitney U test). The phages had a median GC content of 61.17%, while the bifidobacteria harbouring prophages had a median GC content of 59.77%.

The genomic diversity of all 1,076 computationally predicted prophages was explored to find evidence for their mobility and potential for in-host recombination. Pairwise nucleotide identity analysis revealed a high degree of genomic diversity as the majority of prophages shared an intergenomic similarity of below 70% (Figure 4). There were multiple regions of clustering with high levels of intergenomic similarity, which was likely indicative of genus-rank groupings. At peripheral regions of the heatmap, there were several areas of intermediate intergenomic similarity. This observation could be indicative of previous recombination events occurring between distantly related prophages, leading to the formation of “mosaic” genomes. Though, despite the stringent quality thresholds set in our prophage recovery pipeline, these regions of similarity could still be bacterial contamination from incorrectly defined prophages ends. Although some clusters of prophages were exclusively from a single host species, there were also several regions that included groups of closely related phages that were identified across multiple different hosts. For instance, certain clusters showing high intergenomic similarity were from prophages identified in *B. adolescentis*, *B. catenulatum* and *B. pseudocatenulatum* strains. There appeared to be no discernible grouping of phages based on geographic location, age, or sample type. However, much of the metadata in these categories was absent from public databases.

**Figure 4.**
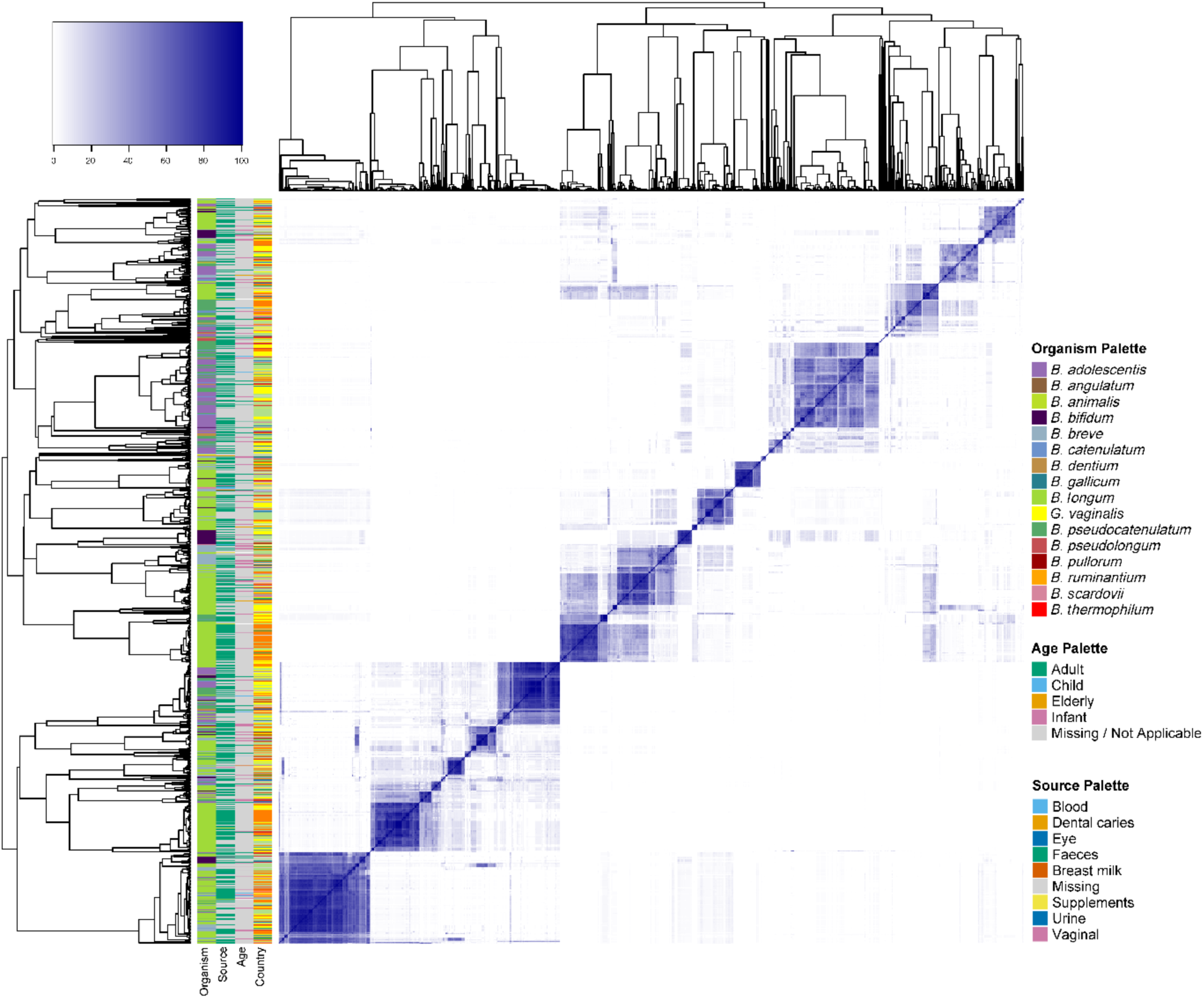
Dendrogram and pairwise comparison heatmap showing intergenomic similarity of 1,076 prophage genomes. There were 22 regions of clustering (blue boxes), which show high intergenomic similarity. The coloured columns (left) represent host species (Organism) and metadata, including isolation source (Source), age group (Age), and country where bacterial host was isolated (Country; see Table S1).

To explore the evolutionary relationships between prophages, a proteomic tree was constructed based on global genomic similarities at the amino acid level (Figure 5). The phages appeared to initially split into two larger groups, which could represent a minimum of two orders. Among these groups, prophages exhibited clustering within several long and short branching clades. The short branches were likely representatives of the same species with recent divergence, while the deeply branched clades were suggestive of phages with distinct evolutionary trajectories. Furthermore, there were multiple regions of the tree where groups of phages within a clade appeared to infect different hosts, suggesting mobilisation of phages within the genus. Prophages from *B. adolescentis*, *B. catenulatum* and *B. pseudocatenulatum* strains appeared to group together in multiple clades, which was indicative of phages that share a more dynamic horizontal mobilisation pattern within these species. For further exploration of their phylogenetic relationships, a viral proteomic tree was also constructed which included all known isolated bacteriophage genomes and *Bifidobacterium* prophage genomes identified in this study (Figure S5). Despite bifidophages demonstrating high levels of genomic diversity, they all cluster into four distinct deeply branched clades, sharing few similarities with reference genomes. These observations signify the presence of at least three novel phage orders. As each clade has long branching roots, this was indicative of early divergence in the evolutionary histories of each group and many expected substitutions in their protein sequences.

**Figure 5.**
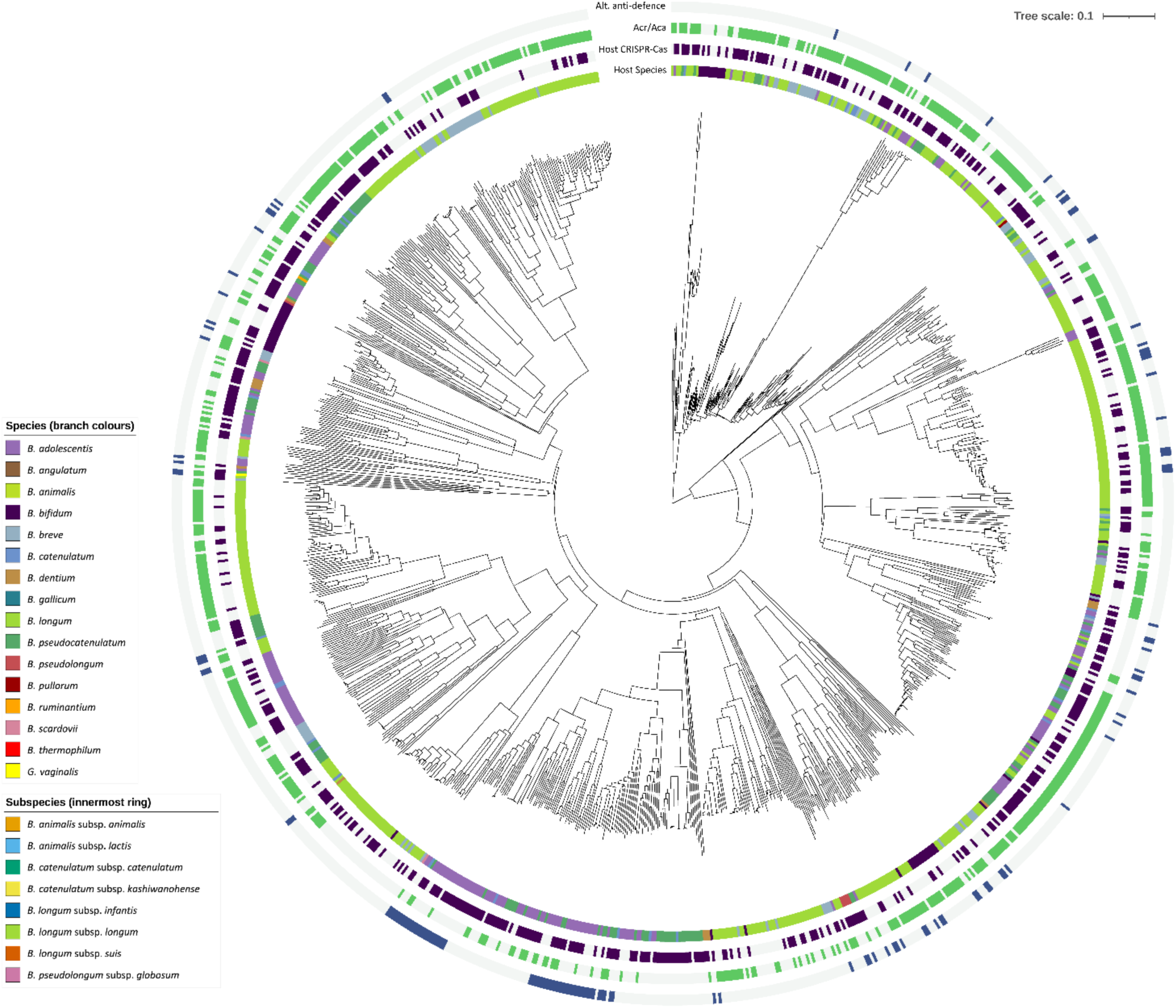
Viral proteomic tree of all computationally predicted prophages included in this study. The inner ring is coloured in accordance with the host species, while the second ring represents presence (purple) or absence (grey) of host CRISPR-Cas. The outermost rings represent the presence of Acr/Aca proteins (green) and alternative anti-defence systems (blue) in prophage genomes. The tree scale and branch lengths correspond to pairwise distance between prophages.

All bifidobacteria-derived proviruses were identified as likely novel *Caudoviricetes* phages, as none could be taxonomically classified into any known genus or species. Closer inspection of their intergenomic differences revealed that there were 278 putative genera and 443 species representatives. Species representatives were selected based on the largest genome length for each cluster.

### *Bifidobacterium* prophages harbour several unique counter-defence genes

To investigate if *Bifidobacterium* prophages have adapted to circumvent the defence systems of their host, we initially aimed to detect anti-CRISPR (Acr) and anti-CRISPR-associated (Aca) proteins in viral sequences. Of 1,076 prophages, 660 encoded at least one Acr/Aca protein (∼61%), with 3,242 genes identified in total (Figure 5). There was a significant association between prophages encoding Acr/Aca proteins with the presence of CRISPR-Cas systems in host strains (Chi-square statistic of 4.17587 and a simulated p-value of <0.05). Though, there was no significant association found between the presence of an Acr/Aca protein and the prophage being targeted by CRISPR spacers (Chi-square statistic of 0.9963391 and a simulated p-value of 0.32).

Alternative anti-defence systems were also identified in 136/1,076 (∼13%) of prophages, with 141 individual counter-mechanisms detected. The most abundant anti-defence system targeted toxin-antitoxin mechanisms (55 genes). There were also counter-mechanism strategies targeting RecBCD (40 genes), RM (20 genes), BREX (2 genes), Gabija (1 gene) and Hachiman (1 gene). There were also 22 genes for broad-spectrum counter defence. Preliminary investigations of alternative proteins that could potentially influence host evolution revealed that 122 bifidophages encoded a WhiB transcriptional factor, which is a regulator found exclusively in Actinobacteria (114). Where prophage-encoded WhiB is basally expressed in mycobacteria, this induces a phage-resistant phenotype in the host (115). Therefore, the presence of these proteins and other undeciphered genes in bifidobacteria-derived prophages may bear influence over their host in the human environment, which warrants further exploration.

## Discussion

In this study, we investigated the hypothesis that arms race dynamics are occurring between bifidobacteria and their prophages in the human environment. To test this, we needed to demonstrate that (i) human-associated bifidobacteria incorporate antiviral defence systems into their genomes that specifically target phages, (ii) bifidobacteria harbour prophages and (iii) prophages counteract the bacterial defence systems by encoding anti-defence systems in their own genomes.

For this investigation, the results from both DefenseFinder (40) and PADLOC (78) were combined when detecting anti-phage defence systems. Although each tool uses MacSyFinder (116), and Hidden Markov Models (HMMs) to screen genomes for the presence of defence systems via homology and genetic architecture comparison, they each screen against different databases. The importance of using both tools was also highlighted in the final dataset of defence systems. Initially, DefenseFinder detected just over 6,000 defence systems, equating to around 6 defence systems per genome. When combining both tools, this figure doubled to an average of ∼12 defence systems being encoded by each strain, with PADLOC identifying several thousand anti-viral mechanisms that DefenseFinder did not. Though, had PADLOC been used on its own, there would have been ∼2,000 fewer systems in the final dataset. This finding highlights the importance of employing a multi-faceted approach in recovering defence systems.

By conducting a systematic analysis of over 1,000 human-associated *Bifidobacterium* genomes, a diverse and complex repertoire of anti-viral defence systems was identified. For this study, the *Bifidobacterium* strains had a median genome size of ∼2.3Mbp, ranging from just ∼1.6Mbp to ∼3.1Mbp, which is less than half the average of 5Mbp for ‘typical’ bacteria (45). Despite their small genome size, the dataset of human-associated *Bifidobacterium* genomes used in this study encoded more than double the number of defence systems than have been reported for other bacteria (117). Here, bifidobacteria were discovered to harbour around 12 defence systems per genome, compared to between five and seven defence systems for bacteria with average genome sizes (5Mbp), such as *E. coli* (40, 43, 58). Though, Georjon *et al*. (2023) only used one defence system detection tool, namely DefenseFinder, to identify antiviral systems (58).

With around five defence systems, “average” bacterial genomes will encode around one defence system per megabase, while human-associated bifidobacteria, will carry one defence system per ∼268Kbp, suggesting a large metabolic dedication to defence from viruses and other invasive mobile genetic elements. This could perhaps be attributed to the ancient evolutionary origins of bacteria belonging to the Actinobacteria phylum (118, 119), with bifidobacteria representing one of the most deeply branched lineages within this bacterial group (120). It is thought that members of the *Bifidobacteriaceae* family have co-evolved with hominids for the last 15 million years (121), suggesting a possible long occurrence of arms race dynamics between bifidobacteria and phages. Furthermore, as bifidobacterial evolution is greatly linked to their host/niche (122, 123), and with their evolutionary trajectory likely undergoing numerous loss and gain events (124), this organism has demonstrated to be highly adaptable and versatile. If their defence systems were not required for survival, bifidobacteria would have likely lost them over time.

Our analysis revealed that certain defence systems in bifidobacteria, such as RM, occurred at similar frequencies when compared to most bacteria (40, 125, 126). The prevalence of Abi and CRISPR-Cas systems was found to be higher in the *Bifidobacterium* genomes here than in most prokaryotes, occurring in ∼83% and ∼59% of strains, respectively, as Abi is reported to occur in ∼73% of all fully sequenced bacterial genomes (126), and CRISPR-Cas is present 42-46% of bacteria (127, 128). The findings for CRISPR-Cas systems align with previous observations of CRISPR-Cas systems in *Bifidobacterium* spp., which found systems in ∼57% of strains (57). However, there were notable differences between the abundance of certain defence systems across individual species. This variation could be a result of several factors, such as different selection pressures based on varied exposure to phages, as well as increased horizontal transfer among strains or species harbouring CRISPR-Cas systems with flanking ABC transporters (129). Additionally, the spacer sequences present in bifidobacteria matched ∼74% of prophages identified in this study. CRISPR arrays found in other bifidobacteria have also been found to specifically target prophages in alternative studies (49, 56, 57). These results provide evidence that the CRISPR-Cas systems in human-associated bifidobacteria specifically target viruses and are capable of actively adapting to and repelling infecting temperate phages and possibly other mobile genetic elements. More specifically, the vast majority of spacers that targeted prophages in this study belonged to Type I CRISPR systems, suggesting that Type I systems primarily target phages while Type II systems may be targeting alternative mobile genetic elements, such as plasmids or these systems may target phages that are currently undiscovered.

Present in over 90% of strains, this dataset of human-associated bifidobacteria exhibited a notably higher prevalence of Wadjet systems than in other Actinomycetota (43%) and bacterial species overall (7%) (58, 130, 131). Wadjet was highly abundant in all species (>5 strain representatives) except *B. breve*, where this defence mechanism was completely absent. As Wadjet has been previously characterised as an anti-plasmid system (131), this may explain the lack of plasmids in bifidobacterial cells (132), and those containing plasmids appear to be highly specialised to the bacterial host (133, 134). The substantially high prevalence of Wadjet within this genus may also be a contributory factor for its reputation as a notoriously difficult organism to transform (135). A previous study found that transplantation of a functioning Wadjet operon into *Bacillus subtilis* significantly reduced the frequency of successful transformations (130). Additionally, *B. breve*, which lacks the presence of Wadjet systems, has been previously shown to be transformable (136, 137). More recently, *B. breve* strain UCC2003 has also been transformed using a barcoded transposon insertion pool (138). If gene knockouts can be successfully introduced in Wadjet systems encoded by bifidobacteria in future, this could perhaps be a considerable step towards overcoming existing difficulties in its genetic manipulation.

As part of the analysis to decipher the defence systems present in bifidobacteria, we also wanted to explore patterns of co-occurring defence systems and their potential roles in enhancing protection from infecting phages. Earlier findings have reported synergistic interactions between pairs of defence systems in bolstering protection against infecting phages, such as Tmn co-opting the domain of ATPase enzymes from Gabija in *E. coli* (43). This study revealed that there were several instances where CRISPR-Cas systems positively co-occurred with RM systems, including CRISPR-Cas subtype I-E and RM Type II-G, which has previously been reported to enhance protection from phages in *Streptococcus thermophilus* (139). It has also been observed in *Staphylococcus aureus,* that restriction digestion of viral genomes can result in spacer acquisition to the CRISPR-Cas Type II-A system from the cleaved site, which can trigger a more dynamic immune response against the infecting phage (140). These results may be an indication that similar synergistic activities could be occurring between CRISPR-Cas and other defence systems in bifidobacteria to provide increased protection from phages.

Furthermore, abortive infection-type defence systems were predominantly shown to positively co-occur with alternative systems, with more than half of co-occurring pairs containing at least one of these systems, including CRISPR-Cas. While synergism between CRISPR-Cas systems and abortive infection systems has previously been observed, this has been mainly linked to Type III and Type VI systems, which have been shown to mediate the activation of non-discriminate nucleases (141, 142, 143, 144). Though, Type I CRISPR-Cas systems have been shown to provide immunity to phages at population level by triggering an ‘altruistic’ abortive infection response when CRISPR-Cas fails to completely rid a cell of viral infection (145). Synergism of defence systems with Abi may be a collective altruistic strategy employed by bifidobacteria to provide population-level immunity by restricting the replication of phages when the initial lines of defence fail (145, 146). However, before any definitive conclusions can be made, lab-based experiments would be required to validate these observations. It should also be considered that *Bifidobacterium* strains have a historical reputation for being difficult to genetically alter (135), presenting potential obstacles in laboratory validation, as highlighted when discussing the importance of Wadjet systems. Overall, these results may be indicative of several defence systems working synergistically in bifidobacteria to enhance protection from phages in human-associated environments, in addition to their stable prophages.

There was at least one prophage present in just over two-thirds of *Bifidobacterium* strains used in this study, which was notably lower than a previous report for human-associated strains (∼82%) (147, 148). Though, this could be attributed to the use of more stringent identification methods being employed here, as only prophage sequences that were predicted to be more than 50% complete, with a minimum of four structural proteins comprising at least 10% of CDS were retained. The threshold of 10% structural CDSs was introduced after we observed a significant number of false positive detections of prophages in the CheckV “high quality” category, a more stringent control. Therefore, our results align with previous observations of prophage content within *Bifidobacterium* strains, ranging from 50-80% of strains (32, 56, 149).

*Bifidobacterium* prophages also exhibited high genomic diversity, often encoding genes to circumvent host defences, suggesting counter defence is a key contributary factor in their genetic diversity. Intergenomic similarity comparisons and observations of groups of closely related phages that infect several different hosts, provided strong evidence to suggest that bifidophages may undergo genetic recombination, which has previously been shown to facilitate host range expansion (150). Further characterisation of *Bifidobacterium* prophages revealed that Acr/Aca proteins were encoded by more than half of the dereplicated genomes. In addition, several alternative “anti-defence” systems were also detected, including anti-RM and broad-spectrum counter-defence proteins. Investigations into the abundance and distribution of CRISPR-Cas resistance proteins in prokaryotic viruses has revealed that Acr/Aca candidates were present in 58% of viral genomes (151), which closely reflects the results observed in this study (∼61%).

## Study Limitations

While robust methods for identifying defence systems in bacteria have been developed in recent years (40, 78), there may still be discovery biases associated with the identification of defence systems in bifidobacteria. As a large portion of *Bifidobacterium* genes remain undeciphered, and with bifidobacteria being highly specialised and adapted to their hosts or environment (122, 152, 153), it is possible that genus-or species-specific defence systems are being missed. Additionally, the vast majority of defence systems that can be identified using DefenseFinder (40) originated from investigation in bacteria belonging to the Pseudomonadota phylum (58). There have also been just three mechanisms originally identified or tested in Actinomycetes (154, 155, 156), with no known systems currently originating from *Bifidobacterium* species. Thus, there may be biases associated with currently available tools, as well as many defence systems yet to be uncovered. These analyses may facilitate the systematic targeting of hypothetical proteins for further characterisation, due to bacterial defence systems tending to co-localise within genomes in so-called “defence islands”. It has recently been reported that defence systems in *E. coli* cluster within mobile island “hotspots” that are flanked by specific core genes (157). By employing a similar methodology, it may be possible to identify unknown genes of interest within these genomic regions to possibly discover novel defence systems, which could be specific to the genus or individual *Bifidobacterium* species. Recently, a ‘guilt-by-embedding’ strategy has been used to identify potential anti-phage candidates and also confirm the function of known defence systems (158).

Despite a rapid expansion in the number of known antiviral defence systems in bacteria (159), there has been a lack of discovery of anti-prokaryotic immune systems (160), with exception of Acr systems (161), where over 100 Acr proteins have been characterised (162). There are also several online databases available containing experimentally verified Acrs (151, 163, 164, 165). This extensive work could explain why anti-CRISPR systems were identified as the most abundant “anti-defence” system in bifidophages. To date, there is just a single bioinformatics resource for the detection of alternative classes of “anti-defence” genes, namely dbAPIS (107). Moreover, likewise to the presence of defence systems in bifidobacteria, there may be similar biases associated with the detection of “anti-defence” proteins. Overall, our study suggests that human-associated bifidobacteria encode numerous defence systems and harbour genomically diverse prophages with counter defence mechanisms.

## Conclusions

This study has highlighted evidence for an “arms race” in the human environment between bifidobacteria and their phages which contributes to genomic diversification of both the bacteria and the phages. *Bifidobacterium* genomes were found to possess a varied arsenal of defence systems that specifically target viral elements, including CRISPR spacers that match almost all three-quarters of predicted prophages. This observation was accompanied by the presence of a genomically diverse pool of prophages that contained several unique anti-immune system genes, including Acr/Aca proteins. We hope that by unravelling these relationships, this research can contribute to the development of high-throughput methods for identifying and testing next-generation live biotherapeutic strains, which can be applied to various fields for the improvement of both human and animal health. Moreover, by deciphering the defence system repertoire of bifidobacteria, this study can facilitate the development of genetic tools for the use of this bacterium in mechanistic studies.

## Data availability and supporting code

Please see BioProject accessions PRJNA958097, PRJNA720750, PRJEB28188 and PRJNA1209344 for further information regarding genome assemblies. A full list of the 1,087 assembly accession numbers used in the final analysis of this study, including accompanying metadata, is available in Table S1 in supplemental material. Supporting code is available at https://sweet-diadem-981.notion.site/Diverse-Defence-Systems-and-Prophages-in-Human-Associated-Bifidobacterium-Reveal-Arms-Race-Dynamic-14bd61222c0380538849facd06996ed9.

## Acknowledgements

We would like to extend thanks to QIB Sequencing Core Services, QIB Bioinformatics Support and NBI Research Computing for their continued assistance. This research was supported in part by the NBI Research Computing through the provision of a high-performance computing cluster. The authors would like to acknowledge the site-specific clinic staff members as well as the mothers and infants for participating in the PEARL study.

This work was supported by the UKRI Biotechnology and BBSRC Doctoral Training Partnership BB/M011216/1. L.J.H. is supported by Wellcome Trust Investigator Award 220876/Z/20/Z; E.M.A. and L.J.H acknowledge funding by the Biotechnology and Biological Sciences Research Council (BBSRC), Institute Strategic Programme Gut Microbes and Health BB/R012490/1, and its constituent projects BBS/E/F/000PR10353 and BBS/E/F/000PR10356. E.M.A. was further supported by the BBSRC Institute Strategic Programme Food Microbiome and Health BB/X011054/1 and its constituent projects BBS/E/F/000PR13631 and BBS/E/F/000PR13633; and by the BBSRC Institute Strategic Programme Microbes and Food Safety BB/X011011/1 and its constituent projects BBS/E/F/000PR13634, BBS/E/F/000PR13635 and BBS/E/F/000PR13636. R.C. and E.M.A. are funded through the Biotechnology and Biological Sciences Research Council (BBSRC) grant Bacteriophages in Gut Health BB/W015706/1.

## Author Contributions

J.A.D.D.: writing, methodology, investigation, data curation, visualisation, formal analysis. E.M.A.: supervision, funding, project administration, conceptualisation, methodology, validation. L.J.H.: supervision, project administration, resources, conceptualisation, methodology, validation. R.C.: supervision, methodology, conceptualisation, validation. T.B.: supervision, conceptualisation. M.K.: resources, methodology, conceptualisation, validation. R.K.: supervision, resources, methodology, conceptualisation, validation. X.D.: methodology, conceptualisation. A.T., S.T.: resources, software. R.S., S.P., R.W.: resources.

## Supplementary Figure Legends

**Figure S1. Global map illustrating the countries of origin for isolated sources.** World map with pie charts that were colour-coded based on the proportions of individual bifidobacterial species. Pie chart size was proportional to the number of representative strains from each country.

**Figure S2. Prevalence of defence system types across the human-associated *Bifidobacterium* genome collection.** Circular bar charts demonstrating the percentage occurrence of defence system types present in at least one *Bifidobacterium* genome.

**Figure S3. Prevalence of defence system types in individual *Bifidobacterium* species.** Circular bar charts demonstrating the percentage occurrence of defence system types present in each *Bifidobacterium* species.

**Figure S4. Tallies of CRISPR-Cas subtypes present in bifidobacteria.** Upset plot showing the number of occurrences of each CRISPR-Cas subtype in human-associated bifidobacteria.

**Figure S5. Viral proteomic tree of filtered RefSeq bacteriophage genomes and the 443 species representatives *Bifidobacterium* prophages identified in this study**. The branches of bifidobacteria-derived prophages were coloured based on their prospective order, which were tentatively named Lughvirales, Dagdaovirales and Morriganvirales. The innermost rings surrounding the tree represent viral families and genera, respectively. The outermost ring (pink) corresponds to bifidophages detected in this study.

## Supplementary Table Legends

**Table S1. Full list of *Bifidobacterium* genomes used in the final analysis of this study.** List of all *Bifidobacterium* strains analysed in this study, including accompanying genomic data, assembly statistics and metadata.

**Table S2. List of updated genome names.** List of 17 genome names that have been updated following phylogenetic analysis and QC checks.

**Table S3. List of defence systems with conflicting calls by DefenseFinder and PADLOC, along with the system included in the final results.**

**Table S4. Matches of *Bifidobacterium*-derived CRISPR spacers with prophage sequences**.

